# metaSMASH: Scalable Biosynthetic Gene Cluster Detection for Large Metagenomic Assemblies

**DOI:** 10.64898/2026.07.22.739405

**Authors:** Caner Bağcı, Kai Blin, Nadine Ziemert

## Abstract

antiSMASH is widely used for biosynthetic gene cluster (BGC) detection and annotation, but its standard workflow is poorly suited to large metagenomic assemblies, where massive contig counts create severe runtime bottlenecks and complicate downstream result exploration. We present metaSMASH, a re-engineered fork of antiSMASH for metagenome-scale BGC analysis. metaSMASH preserves the original antiSMASH detection and annotation logic while introducing streaming, memory-bounded execution, record-level parallelisation, optional output filtering, and an interactive dashboard for large result sets. Across 25 benchmark metagenome datasets, metaSMASH reproduced identical BGC detection results while dramatically reducing computational cost. Relative to the default antiSMASH configuration, metaSMASH was a geometric-mean 38× faster. It also outperformed an *ad hoc* chunked antiSMASH workflow: in the default configuration it achieved a geometric-mean 2.9 × speed-up and 1.7 × lower peak memory, and with extended-analysis modules enabled it was 2.7 × faster and used 3.1 × less memory while completing all datasets, whereas the *ad hoc* workflow ran out of memory on the two largest assemblies. By substantially reducing the computational burden of large-scale metagenome analysis without sacrificing result equivalence, metaSMASH makes routine mining of assembled metagenomes more practical and provides a scalable foundation for natural product discovery from complex microbial communities.

**Graphical Abstract:** 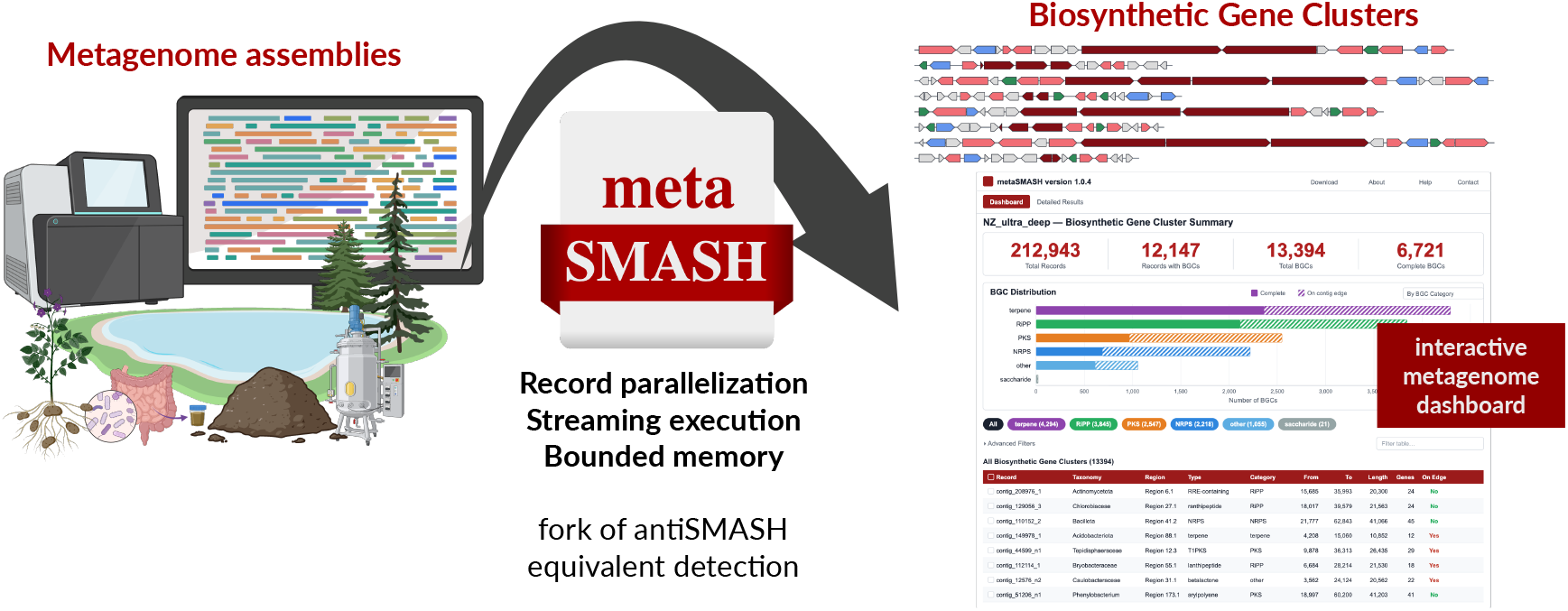

## 1 Introduction

Bacterial natural products are a chemically diverse class of bioactive molecules with important applications in medicine, agriculture, and biotechnology [1, 2]. Because the genes required for their biosynthesis are often co-localized in biosynthetic gene clusters (BGCs) [3], these compounds can be systematically identified and explored directly from genomic sequence data through genome mining. Genome mining has therefore become a powerful strategy for detecting BGCs, prioritizing them for further study, and, in some cases, linking them to known metabolite families [4, 5].

Among the available tools, antiSMASH (antibiotics & Secondary Metabolite Analysis SHell) has become the most widely adopted computational framework for BGC detection and annotation [6]. Since its initial release, antiSMASH has undergone continuous development, and now combines rule-based and profile Hidden Markov Model (pHMM)-based detection with comparative genomics modules such as ClusterBLAST, and rich HTML-based visualization of detected regions. In its current release, antiSMASH 8 can identify BGCs encoding more than 100 classes of secondary metabolites [7].

While antiSMASH performs very well on individual microbial genomes and smaller genome collections, the explosive growth of metagenomic sequencing has created a pressing need for tools that can operate at much larger scales. Modern metagenomic sequencing and assembly workflows routinely generate datasets containing hundreds of thousands to millions of contigs, corresponding to billions of base pairs derived from complex microbial communities [8–10]. These assemblies represent an enormous and still largely untapped reservoir of biosynthetic diversity [10–12], and dedicated resources have begun to catalogue this potential across public metagenomes and genomes [13]. Culture-independent studies across many environments have repeatedly shown that the majority of microbial taxa remain uncultured [10, 14, 15], meaning that much of their metabolic potential can only be accessed through metagenomic approaches.

However, applying antiSMASH to large metagenomic assemblies is severely constrained by its current architecture. The standard antiSMASH pipeline loads all input sequences into memory, converts them into a rich internal data model, and processes them through detection and analysis phases while maintaining all records simultaneously. For large metagenomic assemblies, this design requires the entire dataset to be loaded into memory before analysis can begin, leading to memory demands that can exceed hundreds of gigabytes. In addition, the sequential iteration over records during downstream analysis means that runtime scales poorly with dataset size, often extending to days or weeks for large assemblies. As a consequence, users have had to rely on *ad hoc* workarounds such as splitting inputs into smaller chunks, running multiple parallel instances, and writing custom scripts to merge outputs afterwards. These strategies are difficult to reproduce, prone to error, and do not make efficient use of available computational resources.

Here, we present metaSMASH, a re-engineered fork of antiSMASH designed specifically for metagenome-scale BGC detection and analysis. metaSMASH preserves the full detection and annotation logic of antiSMASH, yielding equivalent BGC predictions while introducing a parallel execution framework with a streaming, memory-bounded architecture tailored to large and fragmented assemblies. In addition, metaSMASH provides incremental output writing, copy-on-write database sharing, optional output filtering, support for taxonomy-aware result exploration, and an interactive HTML dashboard for large metagenomic datasets. Importantly, metaSMASH is distributed as a replacement for antiSMASH and therefore integrates seamlessly with existing databases and downstream workflows, including gene cluster family analysis [16].

## 2 Materials and methods

### 2.1 Software availability

metaSMASH is developed as a fork of antiSMASH and is freely available on GitHub (https://github.com/canerbagci/metasmash), through Bioconda (https://anaconda.org/bioconda/metasmash), and as a Docker container image via BioContainers (https://quay.io/biocontainers/metasmash); an archived version is also deposited on figshare (https://doi.org/10.6084/m9.figshare.33052778). Documentation for the new parameters and modes that differ from upstream antiSMASH is provided on the GitHub repository. The benchmarked metaSMASH release for this manuscript was version 1.0.4, which was built on top of antiSMASH version 8.0.4 with changes up to commit `9e7c145`.

### 2.2 Implementation

metaSMASH extends antiSMASH 8 with several architectural changes that collectively enable scalable processing of metagenomic datasets. All modifications are confined to the orchestration, parallelisation, and output layers; the BGC detection rules, pHMM profiles, and analysis module logic remain identical to the upstream release.

#### 2.2.1 Record-level parallelisation

In the standard antiSMASH pipeline, records (contigs) are processed sequentially, where each contig passes through gene-finding, BGC detection, and analysis before the next contig begins. This process is bounded by the CPU utilization of tools employed by the pipeline, such as hmmscan from the HMMER package [17]. metaSMASH replaces this sequential loop with record-level parallel execution, in which a self-contained worker function encapsulates the entire per-record pipeline, so that independent records can be processed concurrently across a pool of worker processes.

#### 2.2.2 Streaming pipeline and incremental output

For inputs exceeding a small number of records, metaSMASH activates a streaming pipeline that separates BGC detection from detailed analysis into two distinct phases. This separation allows a bounded memory as not all records containing a BGC region are kept in memory until the end of the pipeline.

In Phase 1 (detection), metaSMASH iterates over records in batches. Each batch is dispatched to a pool of N single-threaded workers, and each worker performs gene finding and HMM-based BGC detection on its assigned record. Only the lightweight PFAM [18] caches are pre-loaded before Phase 1, analysis-specific databases (ClusterBLAST, MIBiG, etc.) are deferred to avoid unnecessary memory consumption in detection workers. Records without BGC regions are either serialised directly to the output files, or discarded entirely. Records that contain BGC regions are collected into a bounded window for Phase 2.

Phase 2 (analysis) begins when the bounded window reaches a defined capacity or the input source is exhausted. At this phase, analysis-specific databases are pre-loaded, and the collected records are dispatched to a pool of `W` multi-threaded workers. Each worker runs the full set of enabled analysis modules on its assigned records. As each worker completes, results are written incrementally to output files and the record’s heavy data is stripped to free memory. After the window is drained, the pipeline returns to Phase 1 to fill the next window.

This two-phase streaming pipeline ensures metaSMASH maintains a bounded memory, which can be optimised to number of available CPU cores and dataset size. After each record completes Phase 2, a stripping function removes the memory-heavy DNA sequences, CDS translations, and record-level feature indices from the memory. Only the metadata needed for HTML overview rendering (such as record ID, region boundaries, product types) is retained as a lightweight object.

#### 2.2.3 Output filtering

Metagenomic assemblies typically contain hundreds of thousands to millions of contigs; however, the vast majority of these harbour no BGC regions. Writing all of these empty records to the output files produces unnecessarily large JSON and GenBank files that are slow to parse and transfer, as well as huge HTML outputs that are beyond the capacity of modern browsers.

metaSMASH introduces the`–output-skip-records-without-regions` flag (enabled by default) that omits records without detected BGC regions from the output files. During Phase 1 of the streaming pipeline, records that pass detection without yielding BGC regions are discarded rather than serialised. The identifiers of skipped records are still logged, and the total count of skipped records is reported in the final summary, as well as in the HTML output. For a typical metagenome assembly where fewer than 1% of contigs contain BGC regions, this filter reduces output file sizes by orders of magnitude, and enables interactive visualization of them.

#### 2.2.4 Metagenome dashboard and taxonomy integration

When processing single genomes, antiSMASH produces a single HTML page that provides an overview of all detected BGC regions, and their detailed views. This design works well for inputs with tens of regions, but becomes unresponsive for metagenome outputs that contain thousands of region-containing contigs spread across the assembly.

metaSMASH generates a new metagenome dashboard as the landing page of the HTML output. The dashboard summarizes information across all processed records and presents it in a searchable and filterable table. Each row represents a detected BGC region and displays the contig identifier, region coordinates, predicted product category and type(s), gene count, whether the region is located on a contig edge, and the most similar known cluster from MIBiG [19] along with a similarity percentage. Summary statistics, including the total number of records processed, the number of records with detected BGC regions, total BGC count, and the proportion of complete regions are displayed above the table. Bar charts break down BGC counts by product category and type, distinguishing between complete regions and those on contig edges.

The dashboard optionally integrates taxonomic annotations. Users can supply a tab-separated file (with the`–html-taxonomy option`) mapping contig identifiers to taxonomy lineage strings. When provided, the dashboard displays an additional taxonomy column in the table and enables filtering by taxonomic group. The taxonomy integration enables rapid assessment of which taxa in a metagenome harbour biosynthetic potential, directly within the metaSMASH results viewer.

The per-region detail pages are preserved unchanged from antiSMASH, with the standard region view with gene arrows, domain annotations, ClusterBLAST hits, and all other analysis module outputs; as this is the standard view most users are familiar and comfortable with.

### 2.3 Benchmarking

In order to evaluate the performance of metaSMASH, we assembled a benchmark set of 25 metagenomes spanning a range of assembly sizes, sequencing technologies, and community complexities (Supplementary Table S1): 15 short-read metagenomes of varying size from the SPIRE database [20], five long-read soil metagenomes from the Microflora Danica project [21], four surface-ocean metagenomes (two short-read and two long-read) from a recent high-resolution diel ocean survey [22], and an ultra-deep long-read soil metagenome from the Schönbuch forest [10]. We compared three pipelines:

a single upstream antiSMASH run (referred to as `upstream antiSMASH`), an *ad-hoc* antiSMASH pipeline that processes the input in parallel chunks (referred to as *ad-hoc* antiSMASH), and metaSMASH. All were based on antiSMASH version 8.0.4, and the benchmarked metaSMASH release was version 1.0.4 (built on the same code base, with changes up to commit `9e7c145`). Upstream antiSMASH was run with `–cpus 4`, because antiSMASH does not scale with additional threads and can in fact run more slowly at higher thread counts owing to the overhead of thread creation (data not shown); a separate sweep identified four threads as the best-performing configuration. To keep the comparison fair, the upstream job was nonetheless allocated a full compute node and differed from the other pipelines only in the number of threads it used. Upstream antiSMASH was benchmarked in the default configuration only, as running it with the extended-analysis modules enabled was infeasible owing to its prohibitively high runtime. For the extended-analysis runs of both workflows, we additionally enabled `–clusterhmmer, –tigrfam, –asf, –pfam2go, –tfbs, –cc-mibig, –cb-general, and –cb-knownclusters`. All runs were performed on an HPC Slurm environment, on nodes with two AMD EPYC 7543 (Milan) processors, with 128 threads, 450 GB of RAM, and 7 days of maximum runtime allocated per run. The *ad-hoc* pipeline was built to run parallel jobs on the input data split into 32 chunks, each with 4 threads assigned, using GNU parallel [23]. For each run that completed successfully within 7 days we measured the total walltime, average CPU load, and peak memory usage.

## 3 Results

### 3.1 Detection equivalence and result exploration

All benchmark runs produced exactly identical BGC detection and annotation results between the *ad hoc* antiSMASH workflow and metaSMASH. In every dataset for which both pipelines completed successfully, the same BGC-containing contigs were identified, the same region boundaries and product predictions were reported, and the same downstream annotation outputs were generated. These results demonstrate that the architectural changes introduced in metaSMASH improve scalability without altering the underlying antiSMASH detection and analysis logic, thereby preserving full result equivalence while enabling analysis of much larger metagenomic assemblies.

In addition to reproducing the standard antiSMASH results, metaSMASH provides a metagenome-oriented dashboard that supports exploration of large result sets through a single interactive overview page (Figure 1). The dashboard enables users to browse detected regions across an entire assembly, inspect summary statistics, sort and filter results by product type or region properties, and rapidly identify contigs or BGC classes of particular interest for follow-up analysis. This interface is especially useful for metagenomic datasets, where thousands of detected regions may be distributed across many contigs and samples, making manual navigation through conventional per-record output pages impractical. By combining an overview of dataset-level trends with direct links to standard antiSMASH region pages, the dashboard helps users move efficiently from global exploration to detailed inspection of individual clusters.

**Figure 1:**
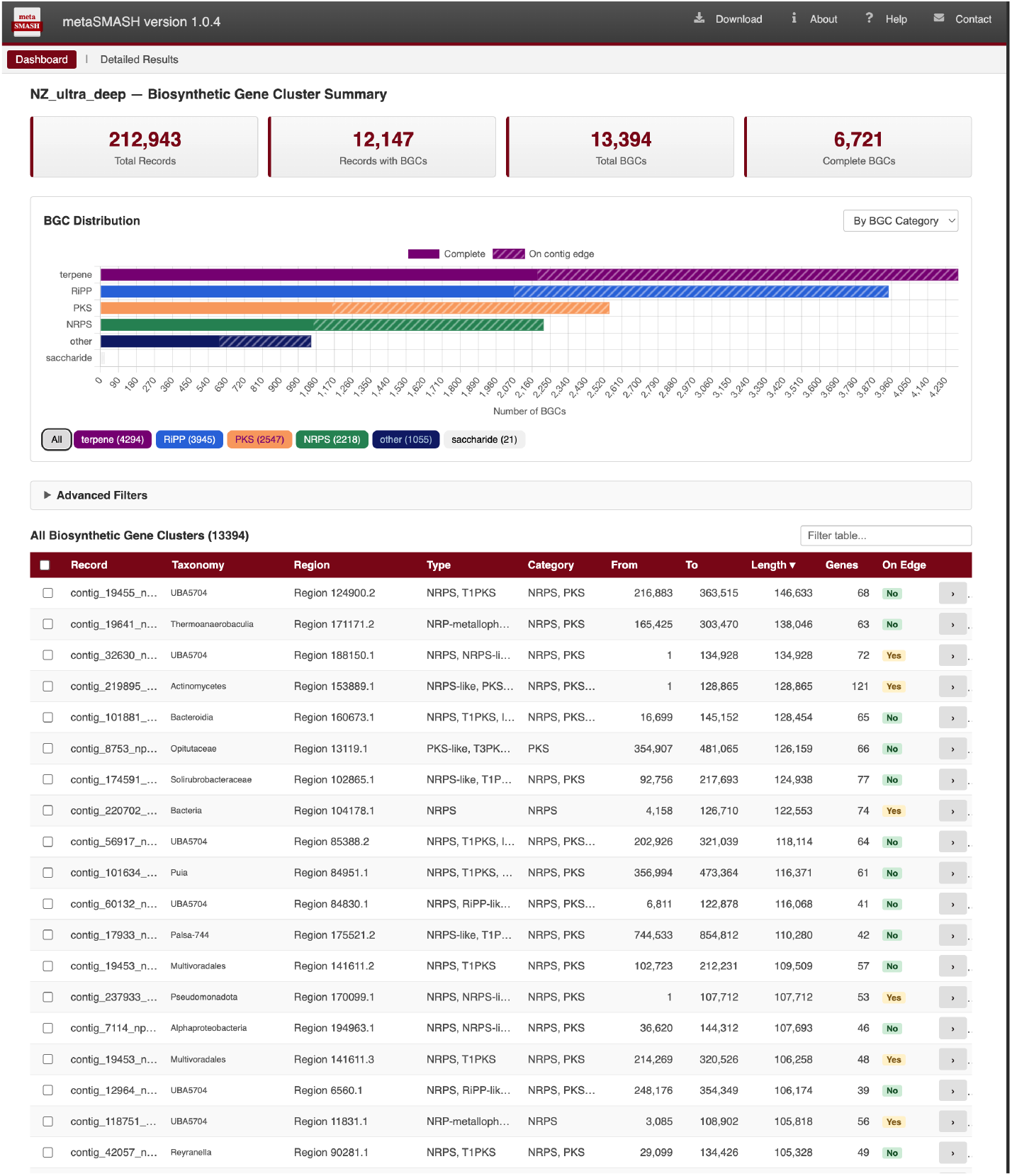
Overview of the metaSMASH interactive dashboard for metagenomic BGC exploration. The dashboard provides a searchable and filterable summary of detected regions across an assembly, including contig identifiers, region coordinates, predicted product classes, gene counts, edge status, and similarity to known reference clusters. Summary panels and charts support rapid inspection of dataset-level patterns, while links to per-region pages enable detailed follow-up analysis of individual BGC predictions.

### 3.2 Runtime performance

Across the 25 benchmark datasets, metaSMASH dramatically outperformed the upstream anti-SMASH configuration, which scales poorly to metagenome-sized inputs. Upstream antiSMASH completed only 24 of the 25 datasets within the runtime limits, failing on one highly fragmented short-read assembly, whereas metaSMASH and the *ad hoc* workflow completed all 25. On the 24 datasets that upstream antiSMASH did finish, metaSMASH was a geometric-mean 38× faster (median runtime 18 min versus 599 min; Supplementary Figure S1).

metaSMASH also outperformed the *ad hoc* chunked antiSMASH workflow, the common workaround of running antiSMASH in parallel over input chunks. It reduced both runtime and peak memory while producing identical detection and annotation results (Figure 2). In the default detection configuration, metaSMASH completed all 25 datasets with a median wall-clock runtime of 22 min versus 47 min for the *ad hoc* workflow (Figure 2A), a geometric-mean speed-up of 2.9 × (per-dataset range 1.1–3.9×). Peak resident memory was likewise lower for metaSMASH on most datasets (median 23 GB versus 42 GB; Figure 2C), a geometric-mean saving of 1.7×.

**Figure 2:**
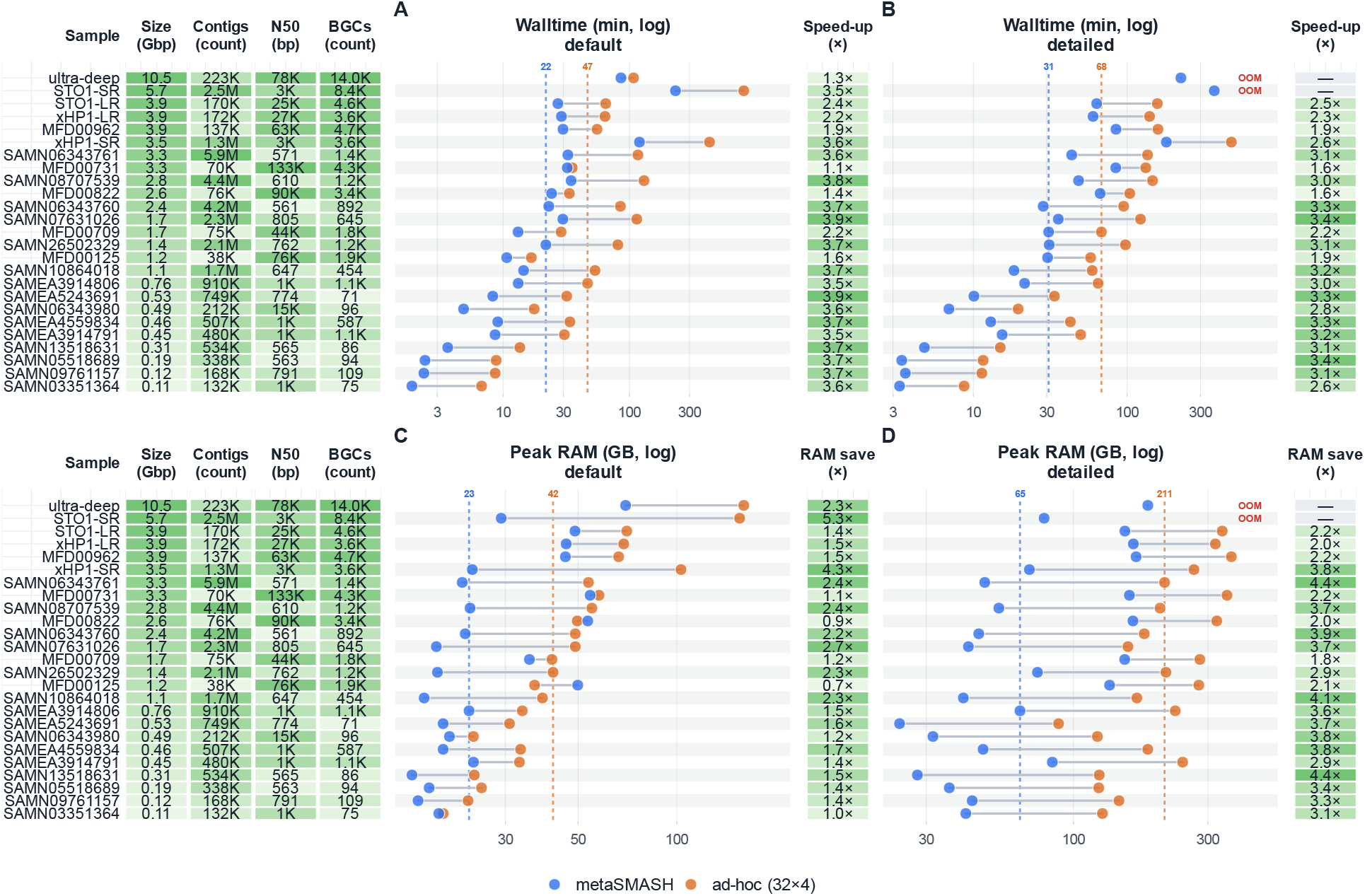
Per-dataset performance comparison of metaSMASH and the *ad hoc* chunked antiSMASH workflow across the benchmark metagenome datasets. Each row corresponds to one dataset, ordered by assembly size; the left-hand tables list dataset characteristics (assembly size in Gbp, number of contigs, contig N50, and number of detected BGCs). (A, B) Wall-clock runtime (minutes, log scale) for the default detection runs (A) and the extended-analysis runs with additional analysis modules enabled (B). (C, D) Peak resident memory (GB, log scale) for the default (C) and extended-analysis (D) runs. In each panel, blue and orange points denote metaSMASH and the *ad hoc* workflow, respectively, joined by a segment for each dataset, and dashed vertical lines mark the across-dataset medians. The adjacent coloured columns report the corresponding per-dataset metaSMASH speed-up (A, B) and peak-memory saving (C, D) relative to the *ad hoc* workflow. “OOM” marks runs in which the *ad hoc* workflow exceeded available memory of 450GB.

The advantage grew when extended-analysis modules were enabled. In this configuration metaSMASH again completed all 25 datasets, whereas the *ad hoc* workflow ran out of memory on the two largest assemblies (the 10.5 Gbp ultra-deep soil metagenome and the 5.7 Gbp STO1-SR ocean metagenome) under the 450 GB memory allocation. Across the datasets that both workflows completed, metaSMASH reduced the median runtime from 68 to 31 min (Figure 2B; geometric-mean 2.7 ×) and, most markedly, lowered peak resident memory from a median of 211 GB to 65 GB (Figure 2D), a geometric-mean saving of 3.1 × . Notably, metaSMASH’s peak memory stayed within a bounded range across the full span of assembly sizes, from the smallest 0.1 Gbp assemblies to the 10.5 Gbp ultra-deep metagenome, remaining well within the 450 GB allocation. This bounded-memory behaviour follows directly from the streaming architecture, which limits the number of region-containing records held in memory at any time, and is what allows metaSMASH to complete large extended-analysis runs that exhaust memory under the chunked workflow. Together, these results show that metaSMASH not only preserves result equivalence, but also makes large-scale metagenomic BGC analysis substantially faster, more memory-efficient, and more robust on the largest assemblies than *ad hoc* parallelisation.

## 4 Discussion

metaSMASH offers a practical advance for large-scale metagenome mining by removing key computational bottlenecks that have limited routine application of antiSMASH to very large metagenome assemblies. By combining streaming execution, bounded memory usage, and record-level parallelisation, metaSMASH makes it feasible to analyse datasets that would otherwise require extensive manual partitioning, repeated job submission, and custom post-processing. This substantially lowers the technical burden on users, improves reproducibility, and enables more systematic exploration of biosynthetic diversity across many samples and environments. Importantly, these gains are achieved without changing the underlying antiSMASH detection and annotation logic, allowing users to benefit from improved scalability while preserving compatibility with established down-stream frameworks and analyses.

A further advantage of metaSMASH is that it improves not only computation, but also usability at metagenomic scale. The dashboard-oriented output provides an efficient way to navigate thousands of detected regions, summarize dataset-level patterns, and prioritize clusters for closer inspection, which is especially valuable for exploratory studies. In more conventional workflows, users often need to manually filter large result sets, cross-reference contig identifiers against separate taxonomy annotations, extract subsets of contigs of interest, and then rerun antiSMASH on those reduced inputs to obtain a manageable and viewable HTML output. This back-and-forth between taxonomy tables, contig-level filtering steps, and antiSMASH region views is cumbersome and error-prone, particularly when many candidate BGCs are considered across samples. Together, these features position metaSMASH as a scalable replacement of antiSMASH for high-throughput metagenomic applications, bridging the gap between trusted BGC detection methodology and the scale required by modern sequencing projects.

### 4.1 Pitfalls and limitations of metagenome BGC mining

Metagenomic BGC mining remains constrained by properties of the underlying assemblies, and these limitations are not resolved by improved computational scalability alone. Metagenomic contigs are often highly fragmented, which can split a single biosynthetic region across multiple sequences, truncate true region boundaries, and increase the proportion of on-contig-edge predictions [24]. This fragmentation complicates interpretation in several ways: one biological BGC may appear as multiple partial regions, raw BGC counts can be inflated, and apparent differences between samples may partly reflect assembly contiguity rather than true differences in biosynthetic potential. Fragmentation also introduces a size-dependent detection bias. Shorter and more compact BGCs are more likely to remain intact on individual contigs, whereas large modular pathways are disproportionately broken apart. In particular, nonribosomal peptide synthetase (NRPS) and polyketide synthase (PKS) clusters [25] are often difficult to recover completely and may be detected only as disconnected fragments or as partial edge-associated predictions [26].

Additional technical factors further influence which BGCs are recovered and how confidently they can be interpreted. Assembly quality, sequencing depth, community complexity, and strain heterogeneity all affect whether a cluster is reconstructed completely enough to be detected and classified [27]. Low-abundance organisms may contribute too little coverage for reliable assembly, repetitive biosynthetic regions may assemble poorly, and microdiversity among closely related strains can either split similar loci across multiple contigs or collapse distinct variants into consensus sequences [28]. Chimeric contigs or local misassemblies may further distort gene order and region structure, while short contigs may not contain enough contextual sequence for reliable boundary definition or taxonomic assignment. As a result, both the number and composition of detected BGCs can be sensitive to upstream sequencing and assembly choices, and their performance.

## 5 Conclusion

In summary, metaSMASH extends antiSMASH to metagenome-scale applications by introducing a streaming, parallel, and memory-bounded architecture without altering the underlying BGC detection and annotation logic. Across benchmark datasets, metaSMASH reproduced identical results while delivering substantial gains in runtime and practical usability, and the addition of a dashboard-oriented interface improves exploration of large collections of predicted regions. These features make metaSMASH a practical solution for routine large-scale mining of assembled metagenomes.

At the same time, computational scalability does not eliminate the fundamental challenges of metagenomic BGC mining, including assembly fragmentation, inflated counts from partial regions, incomplete recovery of large clusters, and uncertainty in linking predicted loci to their products and producing organisms. Nevertheless, by lowering the computational barrier to analysis and improving access to large metagenomic datasets, metaSMASH provides a robust foundation for future natural product discovery efforts from environmental microbiomes.

## Abbreviations

BGC: 
NRPS: 
PKS: 
pHMM: 
MIBiG: 

## 8 Data availability

All sequencing datasets analysed in this study are publicly available. The benchmark metagenome assemblies were drawn from the SPIRE database, the Microflora Danica project, the surface-ocean diel survey of Tucker *et al*., and an ultra-deep Schönbuch soil metagenome.

## 9 Author contributions statement

C.B.: Conceptualization, Methodology, Software, Validation, Formal analysis, Investigation, Data curation, Visualization, Writing – original draft, and Writing – review & editing. K.B.: Methodology, and Writing – review & editing. N.Z.: Supervision, Resources, Funding acquisition, Project administration, Writing – original draft, and Writing – review & editing. All authors read and approved the final manuscript.

## 10 Acknowledgments

The authors acknowledge support by the High Performance and Cloud Computing Group at the Zentrum für Datenverarbeitung of the University of Tübingen, the state of Baden-Württemberg through bwHPC and the German Research Foundation (DFG) for funding under project number 455787709 (bwForCluster BinAC 2). The graphical abstract was created with BioRender.com.

