## Supplementary Material for "metaSMASH: Scalable Biosynthetic Gene Cluster Detection for Large Metagenomic Assemblies"

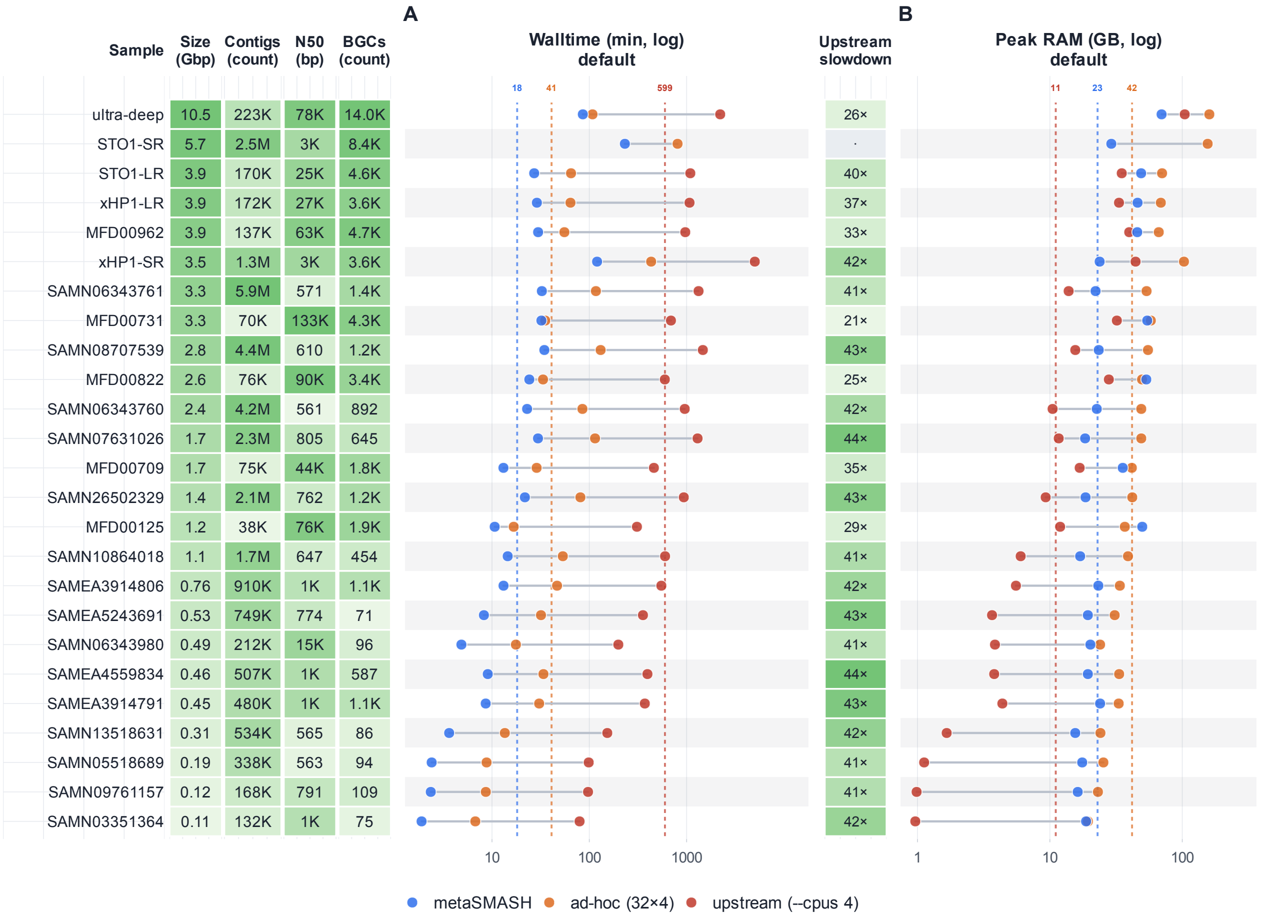


**Figure S1:** Per-dataset performance comparison including upstream antiSMASH, for the default detection configuration. Each row corresponds to one benchmark dataset, ordered by assembly size; the left-hand tables list dataset characteristics (assembly size, contig count, contig N50, and number of detected BGCs). **(A)** Wall-clock runtime (minutes, log scale) and **(B)** peak resident memory (GB, log scale) for metaSMASH (blue), the *ad hoc* chunked antiSMASH workflow (orange, 32×4), and a single upstream antiSMASH instance (red, `--cpus 4`). Points are joined by a segment for each dataset, and dashed vertical lines mark the across-dataset medians. The "Upstream slowdown" column reports the per-dataset runtime of upstream antiSMASH relative to metaSMASH. Upstream antiSMASH did not complete on the largest assemblies within the runtime and memory limits and is therefore shown without a data point (and with a blank slowdown entry) for those datasets.

**Table S1:** Characteristics of the 25 metagenome assemblies used for benchmarking. Datasets are ordered by assembly size. "Reads" indicates the sequencing type; "Size" is the total assembled length; "Contigs" the number of contigs; "N50" the contig N50; and "BGCs" the number of biosynthetic gene cluster regions detected. Datasets were obtained from the SPIRE database (Schmidt *et al.*, 2024), the Microflora Danica project (Singleton *et al.*, 2025), a high-resolution surface-ocean diel survey (Tucker *et al.*, 2025; samples STO1 and xHP1), and an ultra-deep long-read soil metagenome from the Schönbuch forest (Bağcı *et al.*, 2025).

| Dataset | Source | Reads | Size (Gbp) | Contigs | N50 (bp) | BGCs |
|---|---|---|---|---|---|---|
| ultra-deep | Soil (Schönbuch) | Long | 10.47 | 223,441 | 77,794 | 14,008 |
| STO1-SR | Surface ocean | Short | 5.73 | 2,468,893 | 2,504 | 8,432 |
| STO1-LR | Surface ocean | Long | 3.92 | 170,280 | 24,953 | 4,577 |
| xHP1-LR | Surface ocean | Long | 3.91 | 171,698 | 26,878 | 3,618 |
| MFD00962 | Soil (Microflora Danica) | Long | 3.90 | 136,887 | 63,085 | 4,675 |
| xHP1-SR | Surface ocean | Short | 3.48 | 1,339,912 | 3,069 | 3,595 |
| SAMN06343761 | SPIRE | Short | 3.31 | 5,864,565 | 571 | 1,433 |
| MFD00731 | Soil (Microflora Danica) | Long | 3.26 | 70,131 | 132,933 | 4,350 |
| SAMN08707539 | SPIRE | Short | 2.79 | 4,422,430 | 610 | 1,243 |
| MFD00822 | Soil (Microflora Danica) | Long | 2.56 | 76,098 | 90,333 | 3,371 |
| SAMN06343760 | SPIRE | Short | 2.38 | 4,239,798 | 561 | 892 |
| SAMN07631026 | SPIRE | Short | 1.71 | 2,337,646 | 805 | 645 |
| MFD00709 | Soil (Microflora Danica) | Long | 1.68 | 74,729 | 43,583 | 1,819 |
| SAMN26502329 | SPIRE | Short | 1.44 | 2,121,387 | 762 | 1,248 |
| MFD00125 | Soil (Microflora Danica) | Long | 1.24 | 37,687 | 76,456 | 1,870 |
| SAMN10864018 | SPIRE | Short | 1.05 | 1,675,501 | 647 | 454 |
| SAMEA3914806 | SPIRE | Short | 0.76 | 909,919 | 1,000 | 1,141 |
| SAMEA5243691 | SPIRE | Short | 0.53 | 749,118 | 774 | 71 |
| SAMN06343980 | SPIRE | Short | 0.49 | 211,578 | 14,507 | 96 |
| SAMEA4559834 | SPIRE | Short | 0.46 | 507,068 | 1,143 | 587 |
| SAMEA3914791 | SPIRE | Short | 0.45 | 479,729 | 1,334 | 1,077 |
| SAMN13518631 | SPIRE | Short | 0.31 | 534,009 | 565 | 86 |
| SAMN05518689 | SPIRE | Short | 0.19 | 338,368 | 563 | 94 |
| SAMN09761157 | SPIRE | Short | 0.12 | 168,036 | 791 | 109 |
| SAMN03351364 | SPIRE | Short | 0.11 | 132,329 | 1,028 | 75 |
